# ProkSeq for complete analysis of RNA-seq data from prokaryotes

**DOI:** 10.1101/2020.06.09.135822

**Authors:** A K M Firoj Mahmud, Soumyadeep Nandi, Maria Fällman

**Affiliations:** Laboratory for Molecular Infection Medicine Sweden (MIMS), Umeå Centre for Microbial Research (UCMR), Department of Molecular Biology, Umeå University, SE-901 87, Umeå, Sweden; Amity Institute of Integrative Sciences and Health, Amity University Haryana, Gurgaon, 122413, Haryana, India

## Abstract

**Summary:** Since its introduction, RNA-seq technology has been used extensively in studies of pathogenic bacteria to identify and quantify differences in gene expression across multiple samples from bacteria exposed to different conditions. With some exceptions, the current tools for assessing gene expression have been designed around the structures of eukaryotic genes. There are a few stand-alone tools designed for prokaryotes, and they require improvement. A well-defined pipeline for prokaryotes that includes all the necessary tools for quality control, determination of differential gene expression, downstream pathway analysis, and normalization of data collected in extreme biological conditions is still lacking. Here we describe ProkSeq, a user-friendly, fully automated RNA-seq data analysis pipeline designed for prokaryotes. ProkSeq provides a wide variety of options for analysing differential expression, normalizing expression data, and visualizing data and results, and it produces publication-quality figures.

**Availability and implementation:** ProkSeq is implemented in Python and is published under the ISC open source license. The tool and a detailed user manual are hosted at Docker: https://hub.docker.com/repository/docker/snandids/prokseq-v2.1, Anaconda: https://anaconda.org/snandiDS/prokseq; Github: https://github.com/snandiDS/prokseq.

## Motivation

The advancement of massive parallel sequencing and dramatic reduction in sequencing costs have made deep sequencing of RNA (RNA-seq) a primary tool for identifying and quantifying RNA transcripts. Today RNA-seq is widely used to analyse bacterial gene expression in studies that aim to identify drug targets, predict novel gene regulatory mechanisms, etc. Such studies often require profound knowledge of both computational data handling and biology. There are some stand-alone pipelines and tools that require only moderate knowledge of bioinformatics (Delhomme, et al., 2012; Prieto and Barrios, 2019), but these are not designed for analyses of bacterial gene expression.

Prokaryotic RNA-seq analysis is challenging because most available RNA-seq packages assume the input data reflect eukaryotic gene structures, which in many aspects differ from those of prokaryotes (Johnson, et al., 2016). Bacterial transcripts do not have introns and are not alternatively spliced; therefore, using an aligner developed to consider splice junctions often increases falsely assigned reads in the genome (Magoc, et al., 2013). Moreover, unlike in eukaryotes, under specific stresses the expression of almost all prokaryotic genes can change (Creecy and Conway, 2015). Furthermore, quality trimming, adapter removal, and normalization of skewed data are often required for prokaryotic data due to variations in experimental setups, the presence and overexpression of plasmid genes, and differences in RNA-seq protocols (Magoc, et al., 2013; McClure, et al., 2013).

Although there are a few software packages available for prokaryotes that can facilitate the analysis of RNA-seq data, such as SPARTA (Johnson, et al., 2016), EdgePRO (Magoc, et al., 2013), and RockHopper (McClure, et al., 2013), all require substantial knowledge of data handling. Therefore, to reduce human intervention in conducting RNA-seq data analysis for prokaryotes, we developed ProkSeq, a fully automated command-line based workflow. ProkSeq integrates various available tools and built-in functions written in Python. ProkSeq processes RNA-seq data from quality control steps to pathway enrichment analysis of differentially expressed genes. It provides a wide variety of options for differential expression, normalized expression, and visualization, and produces publication-quality figures. Reduced human intervention makes the use of ProkSeq less time consuming than the sequential application of separate tools, which often requires reformatting data. In addition to the convenience, the multithreading feature of the ProkSeq makes the pipeline less time-consuming.

## Implementation

ProkSeq runs in a Linux-based command-line environment and depends on user-defined parameters and sample files. Since it integrates several tools, default parameters for the packages are set in the parameter file. However, the user has the flexibility to change the settings in the parameter file for any desired analysis. The sample file provides the names of the fastq files to be included in the analysis, and also defines the experimental classes, such as treatment and control samples. ProkSeq first checks the quality of reads and filters out low-quality reads using FastQC (http://www.bioinformatics.babraham.ac.uk/projects/fastqc) and afterQC (Chen, et al., 2017). It maps the reads to the reference genome using bowtie2 (Langmead and Salzberg, 2012) and its default parameters for both single and paired-end reads. ProkSeq then generates a report on the alignment quality for each library, as both figures and text, providing information about coverage uniformity, distribution along protein coding sequences, and 5’ and 3’ UTR regions, as well as the read duplication rate and strand specificity generated by RseQC (Wang, et al., 2012) and other built-in functions. Total reads per gene are calculated with featureCounts (Liao, et al., 2014), which provides a high efficiency of read assignments across the genome. ProkSeq also calculates normalized gene expression values for each gene, in the form of transcripts per million (TPM) (Wagner, et al., 2012) and counts per million (CPM) (Wagner, et al., 2012). The formulas by which these are calculated are explained in the supplementary methods (S1). Furthermore, we have integrated salmon (Patro, et al., 2017) to quantify the expression of transcripts by use of a bias-aware algorithm that substantially improves the accuracy and the reliability of subsequent analysis of differential expression.

ProkSeq integrates several tools for differential expression analysis, such as DEseq2 (Love, et al., 2014), edgeR (Robinson, et al., 2010), and NOISeq (Tarazona, et al., 2015). For downstream analysis of differentially expressed genes, ProkSeq uses GO enrichment, and pathway enrichment by integrating clusterProfiler (Yu, et al., 2012). Reports on pre- and post-alignment quality statistics and graphical visualization are created in pdf and HTML formats. One important unique feature of ProkSeq is the integration of well-established normalized methods for skewed data (Creecy and Conway, 2015; Zhu, et al., 2019). Furthermore, the package generates a single-nucleotide resolution wiggle file for visualization in any genome browser. ProkSeq generates vibrant graphics and publication-ready figures at every step of data analysis to give the user more confidence in and understanding of their data. The methods are described in detail in the supplementary methods (S1).

## Sample data sets and results

ProkSeq is distributed with an example data sets. The data set contains paired-end reads from *Yersinia pseudotuberculosis YPIII* (data unpublished) and compares the control versus bile treated. The following files are bundled with exampleFiles.tar.gz.

1. Sample files (sampleCtrl_1.R1/R2.fq, sampleCtrl_2.R1/R2.fq, sampleCtrl_3.R1/R2.fq, sampleTreat_1.R1/R2.fq, sampleTreat_2.R1/R2.fq, sampleTreat_3.R1/R2.fq),
2. Sample description files (samples.bowtie.PEsample, samples.bowtie.SEsample, samples.salmon.PEsample, samples.salmon.Sesample),
3. Parameter definition files (param.input.bowtie and param.input.salmon)
4. Annotation files (oldAnnotationGFF.bed, oldAnnotationGFF.gtf),
5. Transcript file (orf_coding_all.fasta), and
6. Genome file (SequenceChromosome.fasta).

To run the pipeline, one can follow the instructions in https://github.com/snandiDS/prokseq/blob/master/README.md.

We strongly recommend using Docker to run the pipeline. However, the implementation of ProkSeq is straightforward.

1. mkdir prokseq
2. cd prokseq Download the package from github or install using conda.
3. Untar the depend.tar – Depend folder contains all the required external packages. Most binaries of the packages are stored in this folder.
4. Untar the exampleFiles.tar.gz
5. Install the following R packages:

1. DESeq2
2. edgeR
3. NOISeq
4. clusterProfiler
5. apeglm
6. ggplot2

Once all the dependencies and R packages are in place, and the example files are untared, change the PATH in parameter definition file. The PATH variable in the parameter definition file should point to the packages.

For example:

If the directory ‘prokseq’ is created in /home/user/ in Step 1, and the pypy package is in depend folder inside /home/user/prokseq/, then specify the path as below for all the packages in the parameter definition file.

\# Specify the path to pypy required for running afterqc
PATH PYPY /home/user/prokseq/depend/pypy2.7-v7.2.0-linux64/bin

The following syntax can be used to run ProkSeq

Usage: pipeline-v2.8.py [options] arg Options:

-h, --help= Show this help message
-s SAMPLE_FILE_NAME, --sample=SAMPLE_FILE_NAME
-p PARAMETER_FILE_NAME, --param=PARAMETER_FILE_NAME
-n NUMBER OF PROCESSORS, --numproc=NUMBER OF PROCESSORS

The script is run with PE (paired-end) samples described in samples.bowtie.PEsample, and with the parameters defined in param.input.bowtie. The program is submitted with four processors. The program first check if all the packages are available, and generated a report on the screen. The users can decide to proceed based on the report. The result of the pipeline is stored in the Output folder. The Output folder contains all the alignment file, statistics, pre and post filter QC report as well as result folder. The pipeline with the example data produce plots for various steps some of which are shown in figure 2. Details can be found in the github depository of ProSeq where a sample run folder has been included.

**Figure 1:**
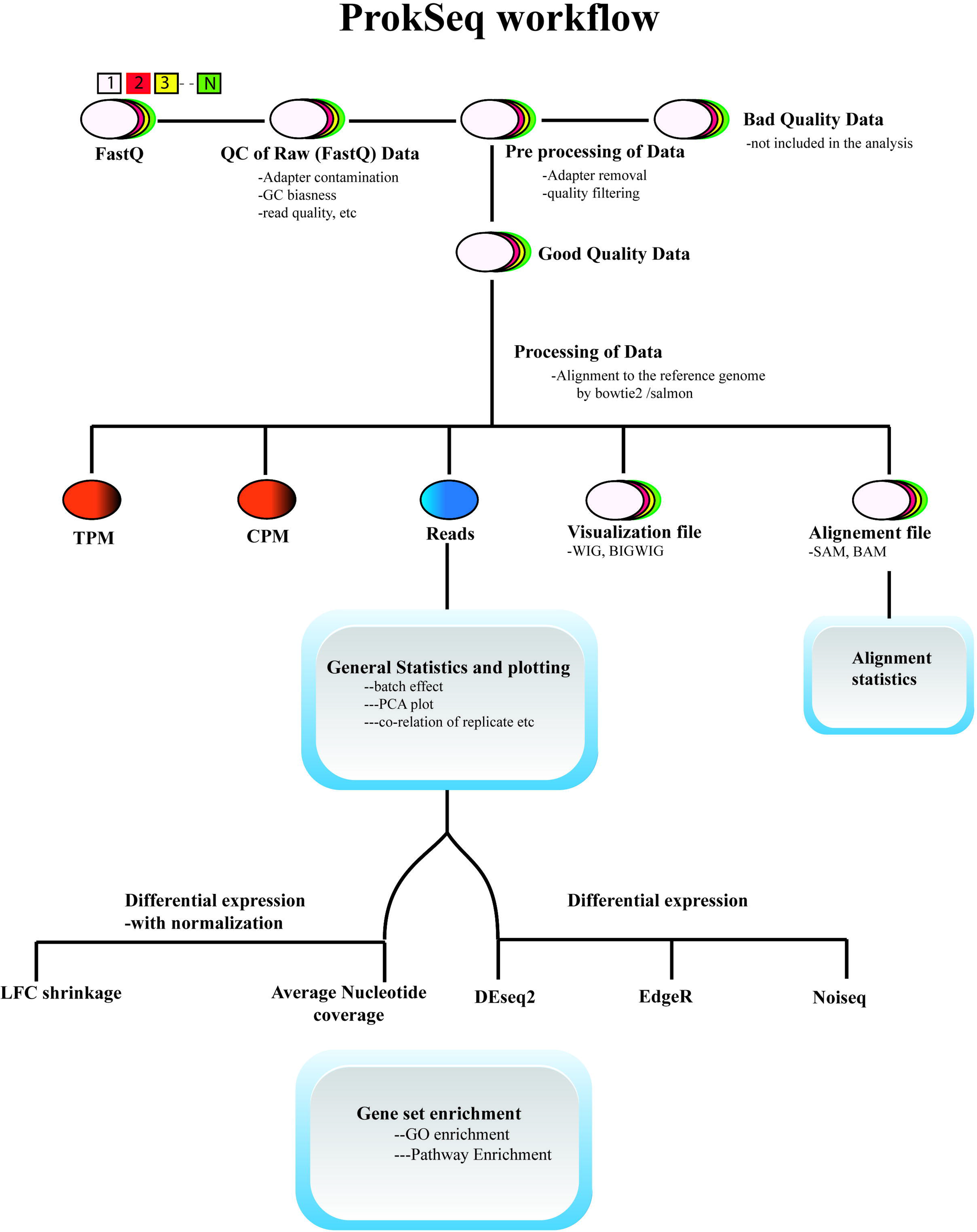
Workflow of ProkSeq showing major steps and tools

**Figure 2:**
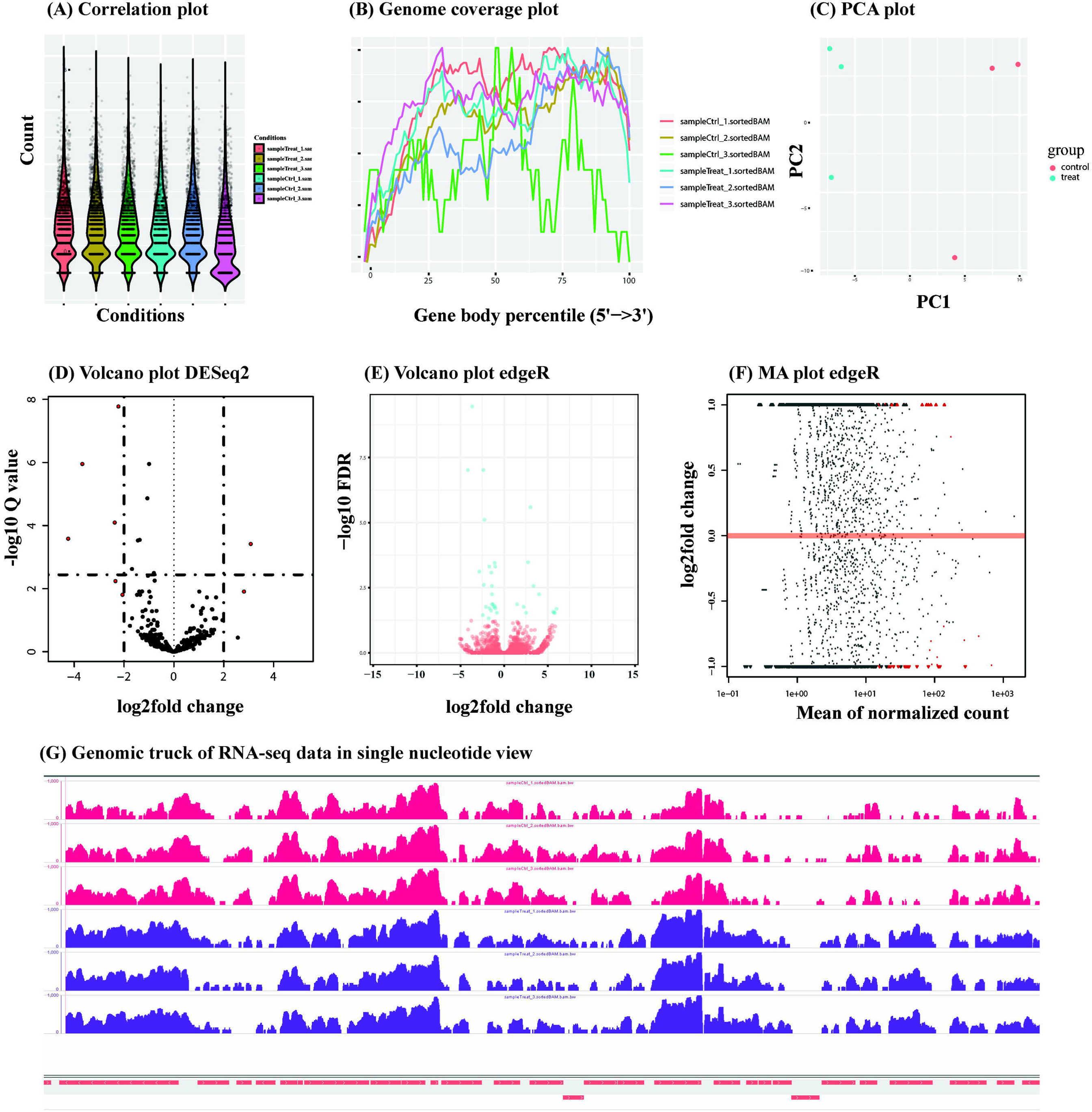
Some of the plots generated by ProkSeq with the sample data sets. (A) Correlation plot showing how the biological replicate and the samples reads are correlated. (B) Genome coverage plot showing how the RNA-seq reads are distributed across the gene. (C) PCA plot of the samples after read count. (D), (E) and (F) Volcano plot and MA plot after differential expression analysis. (G) Single nucleotide genomic track of RNA-seq data is visualized by genomic browser IGV. Single nucleotide genomic coverage is created in the form of wiggle and big wiggle format which can be found in the Output folder.

## Discussion

ProkSeq, which is designed to be used by biologists without significant competence in bioinformatics, will provide new opportunities to discover unique events in transcription dynamics. RNA-seq data can provide much more information than simply the differential expression of known coding sequences. Exploring RNA-seq reads to single-nucleotide resolution across the genome can provide information about biological events other than gene expression. ProkSeq offers easy access to genome-wide visualization of RNA-seq data. Visualization of read mapping will reveal expression from unannotated genomic regions and intergenic regions, including 5’ and 3’ UTRs, which is of great interest in relation to novel transcriptional and translational regulation. Other tools for revealing this type of information that are available today (Table 1) usually require substantial competence in bioinformatics and provide only some of the options available in ProkSeq. Furthermore, integration of salmon in the process gives the user one of the most up-to-date methods of estimating transcript abundance. Salmon uses a realistic model of RNA-seq data that takes into account not only experimental attributes but also biases commonly observed in RNA-seq data. Users can quickly extract transcript abundance and subsequent differential expression data by opting to use salmon.

**Table 1:**
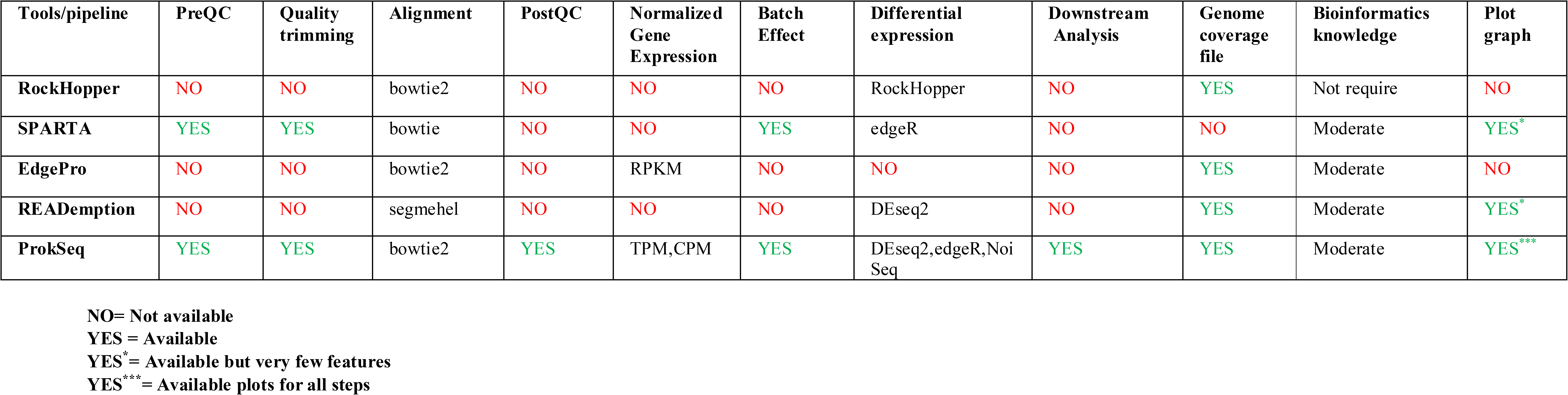
Comparative features of Tools commonly used for Bacterial RNA-seq data analysis

ProkSeq provides an option for batch effect identification with normalization. An essential difference between eukaryotes and prokaryotes that can cause problems when analysing prokaryotic gene expression using tools optimized for analyses of eukaryotic cells is the relative number of differentially expressed genes. Tools such as DEseq2, edgeR, and Limma (Dillies, et al., 2013) are designed with the assumption that most genes are invariant, which is the case in eukaryotes. But in prokaryotes, the expression of the majority of genes is altered under specific stress conditions (Berghoff, et al., 2017; Creecy and Conway, 2015). To address this biasness, ProkSeq normalizes the data at the level of nucleotide base count making the data comparable across samples. ProkSeq provides two normalization options that can handle differential expression analyses of this type of data which are described in detail in the supplementary methods (S1).

ProkSeq has been designed to meet biologists need to analyse RNA-seq data in a reliable and time-efficient way. The built-in automatic sequential handling of the data from differential gene expression analysis to downstream functional analyses allows researchers to focus on complex biological mechanisms instead of tackling bioinformatics obstacles. The flexibility that comes with built-in options for certain steps and the visualisation of mapped reads across genomes opens a path to new discoveries in gene regulation as well as in RNA biology.

## Acknowledgement

The authors thanks Rikki Frederiksen and Chayan Kumar for testing and feed-back, and Dr Nicolas DelHomme from Umea Plant Science Centre for critical reading. The work has been supported by funding from Knut and Alice Wallenberg foundation (2016.0063), Swedish research Council (2018-02855), and the Medical faculty at Umea University.

## References

Berghoff, B.A., et al. RNA-sequence data normalization through in silico prediction of reference genes: the bacterial response to DNA damage as case study. BioData Min 2017;10:30.

Chen, S., et al. AfterQC: automatic filtering, trimming, error removing and quality control for fastq data. BMC Bioinformatics 2017;18(Suppl 3):80.

Creecy, J.P. and Conway, T. Quantitative bacterial transcriptomics with RNA-seq. Curr Opin Microbiol 2015;23:133–140.

Delhomme, N., et al. easyRNASeq: a bioconductor package for processing RNA-Seq data. Bioinformatics 2012;28(19):2532–2533.

Dillies, M.A., et al. A comprehensive evaluation of normalization methods for Illumina high-throughput RNA sequencing data analysis. Brief Bioinform 2013;14(6):671–683.

Johnson, B.K., et al. SPARTA: Simple Program for Automated reference-based bacterial RNA-seq Transcriptome Analysis. BMC Bioinformatics 2016;17:66.

Langmead, B. and Salzberg, S.L. Fast gapped-read alignment with Bowtie 2. Nat Methods 2012;9(4):357–359.

Liao, Y., Smyth, G.K. and Shi, W. featureCounts: an efficient general purpose program for assigning sequence reads to genomic features. Bioinformatics 2014;30(7):923–930.

Love, M.I., Huber, W. and Anders, S. Moderated estimation of fold change and dispersion for RNA-seq data with DESeq2. Genome Biol 2014;15(12):550.

Magoc, T., Wood, D. and Salzberg, S.L. EDGE-pro: Estimated Degree of Gene Expression in Prokaryotic Genomes. Evol Bioinform Online 2013;9:127–136.

McClure, R., et al. Computational analysis of bacterial RNA-Seq data. Nucleic Acids Res 2013;41(14):e140.

Patro, R., et al. Salmon provides fast and bias-aware quantification of transcript expression. Nature Methods 2017;14(4):417–419.

Prieto, C. and Barrios, D. RaNA-Seq: Interactive RNA-Seq analysis from FASTQ files to functional analysis. Bioinformatics 2019.

Robinson, M.D., McCarthy, D.J. and Smyth, G.K. edgeR: a Bioconductor package for differential expression analysis of digital gene expression data. Bioinformatics 2010;26(1):139–140.

Tarazona, S., et al. Data quality aware analysis of differential expression in RNA-seq with NOISeq R/Bioc package. Nucleic Acids Res 2015;43(21):e140.

Wagner, G.P., Kin, K. and Lynch, V.J. Measurement of mRNA abundance using RNA-seq data: RPKM measure is inconsistent among samples. Theory Biosci 2012;131(4):281–285.

Wang, L., Wang, S. and Li, W. RSeQC: quality control of RNA-seq experiments. Bioinformatics 2012;28(16):2184–2185.

Yu, G., et al. clusterProfiler: an R package for comparing biological themes among gene clusters. OMICS 2012;16(5):284–287.

Zhu, A., Ibrahim, J.G. and Love, M.I. Heavy-tailed prior distributions for sequence count data: removing the noise and preserving large differences. Bioinformatics 2019;35(12):2084–2092.

