## Supplementary methods for "ProkSeq for complete analysis of RNA-seq data from prokaryotes": Supplementary methodsS1_tableS2.docx

**Supplementary Methods S1**

**Quality control of reads and filtering:**

Quality control of pre-aligned data is very important for reducing noise in RNA-seq data and getting biologically insightful results. Here we integrated the widely used fastQ quality cheeking tools FastQC (http://www.bioinformatics.babraham.ac.uk/projects/fastqc) and afterQC (Chen, et al., 2017), which can filter out poor-quality data. Because afterQC implements trimming globally by allowing all the reads to be trimmed identically, we have included it as a default setting in ProkSeq so that inexperienced users do not have to think about trimming criteria and bias. In this step, poor-quality reads are eventually filtered out and contaminating adapter sequences are removed. After alignment to the reference genome it is important to get an overview of some of the information in order to interpret the data correctly. RseQC (Wang, et al., 2012) and some built-in functions have been integrated to generate a quality report for each library after alignment

**Mapping to the reference genome:**

ProkSeq maps the reads to the reference genome using bowtie2 (Langmead and Salzberg, 2012) with its default parameters for both single and paired-end reads. Users have the flexibility to change the parameters by changing the parameter.txt file. Bowtie2 was chosen for read mapping due to its flexibility, sensitivity, and lower rate of false mapping (Magoc, et al., 2013) compared with other available short-read aligners used to map bacterial genomes, such as segemehl (Hoffmann, et al., 2009), BBmap (Roy and Chanfreau, 2020), etc.

**Calculated read count and expression per gene:**

ProkSeq calculates total reads per gene with featureCounts (Liao, et al., 2014), which was chosen for its short running times and high efficiency in assigning reads to different genomic features. Most differential expression analysis methods are based on the total number of reads in genes but in some cases when user wants to if the genes are highly expressed or not among the experiment, it is useful to apply normalized gene expression values. For this reason ProkSeq has been designed to calculate counts per million (CPM) (Wagner, et al., 2012) and transcripts per million (TPM)(Wagner, et al., 2012). The formulas by which CPM and TPM are calculated are as follows:

CPM = (number of reads mapped to a gene / total number of mapped reads of a sample) × 10^6^

TPM= (number of reads mapped to a gene/ gene length) / ∑ (reads mapped to a gene/gene length) × 10^6^

**Differential expression analysis:**

It has been demonstrated in several studies that assumptions of equal means and variances do not adequately accommodate RNA-seq data, which are over-dispersed and exhibit great variability between biological replicates (Anders and Huber, 2010). To improve the predictability of differential gene expression, several tools and methodologies have been developed. The most commonly used tools for differential expression analyses are DEseq2 and edgeR, which can all predict differential expression reliably with a limited number of biological replicates. These DE tools mentioned as well as NOISeq are integrated into ProkSeq, the latter being useful for samples with fewer biological replicates (Tarazona, et al., 2015).

**Normalization and differential expression for skewed data:**

Commonly used differential expression methods assume that the numbers of up- and down-regulated genes are balanced between different conditions (Berghoff, et al., 2017). Therefore, these methods might perform poorly when the expression of up to one-fourth of genes is differentially expressed in bacteria in certain biological conditions (Khil and Camerini-Otero, 2002; Kroger, et al., 2013; Yang, et al., 2018). As a consequence, there is a risk that some genes will be wrongly predicted to be differentially expressed. To avoid this, two other normalization methods are included in ProkSeq. These are well-established methods used for analysing bacterial gene expression: (1) gene-specific read alignments, in which each value is expressed as a base count per billion bases counted (Creecy and Conway, 2015). (2) Shrunken log2 foldchanges (LFC) that uses information from all genes to generate more accurate estimates (Zhu, et al., 2019). Normally, lowly expressed genes tend to have relatively higher levels of variability and and LFC shrinkage can reduce it (Zhu, et al., 2019). In the most recent DESeq2 version (>1.16), the shrinkage of LFC estimates is not performed by default. ProkSeq performed it by default and save the result in a file so users can compare with or without LFC shrinkage. However, there is another well-established method that normalize data through in silico prediction of reference genes (Berghoff, et al., 2017). This methods may be included in ProkSeq in the future.

**Data visualization and plots:**

To obtain a view of transcriptome dynamics as a whole, and especially hints about regulatory events such as ribo-switches, small RNAs, and transcriptional start sites, data visualization can be helpful. In addition, visualization also allows scanning of whole genomes to reveal read distributions, which can indicate the expression of intergenic regions and 5´ or 3´ untranslated regions, and alternative transcriptional start sites. ProkSeq provides a default option for data visualization in the form of bigwig files, as well as figures in pdf formats at different steps in the data handling process (Supplementary fig 1).

**References**

Anders, S. and Huber, W. Differential expression analysis for sequence count data. Genome Biol 2010;11(10):R106.

Berghoff, B.A., et al. RNA-sequence data normalization through in silico prediction of reference genes: the bacterial response to DNA damage as case study. BioData Min 2017;10:30.

Chen, S., et al. AfterQC: automatic filtering, trimming, error removing and quality control for fastq data. BMC Bioinformatics 2017;18(Suppl 3):80.

Creecy, J.P. and Conway, T. Quantitative bacterial transcriptomics with RNA-seq. Curr Opin Microbiol 2015;23:133-140.

Hoffmann, S., et al. Fast mapping of short sequences with mismatches, insertions and deletions using index structures. PLoS Comput. Biol. 2009;5(9):e1000502.

Khil, P.P. and Camerini-Otero, R.D. Over 1000 genes are involved in the DNA damage response of Escherichia coli. Molecular Microbiology 2002;44(1):89-105.

Kroger, C., et al. An infection-relevant transcriptomic compendium for Salmonella enterica Serovar Typhimurium. Cell Host Microbe 2013;14(6):683-695.

Langmead, B. and Salzberg, S.L. Fast gapped-read alignment with Bowtie 2. Nat Methods 2012;9(4):357-359.

Liao, Y., Smyth, G.K. and Shi, W. featureCounts: an efficient general purpose program for assigning sequence reads to genomic features. Bioinformatics 2014;30(7):923-930.

Magoc, T., Wood, D. and Salzberg, S.L. EDGE-pro: Estimated Degree of Gene Expression in Prokaryotic Genomes. Evol Bioinform Online 2013;9:127-136.

Roy, K.R. and Chanfreau, G.F. Robust mapping of polyadenylated and non-polyadenylated RNA 3' ends at nucleotide resolution by 3'-end sequencing. Methods 2020;176:4-13.

Tarazona, S., et al. Data quality aware analysis of differential expression in RNA-seq with NOISeq R/Bioc package. Nucleic Acids Res 2015;43(21):e140.

Wagner, G.P., Kin, K. and Lynch, V.J. Measurement of mRNA abundance using RNA-seq data: RPKM measure is inconsistent among samples. Theory Biosci 2012;131(4):281-285.

Wang, L., Wang, S. and Li, W. RSeQC: quality control of RNA-seq experiments. Bioinformatics 2012;28(16):2184-2185.

Yang, B., et al. Global transcriptional regulation by BirA in enterohemorrhagic Escherichia coli O157:H7. Future Microbiol 2018;13:757-769.

Zhu, A., Ibrahim, J.G. and Love, M.I. Heavy-tailed prior distributions for sequence count data: removing the noise and preserving large differences. Bioinformatics 2019;35(12):2084-2092.
